# Nanobody-Functionalized Cellulose for Capturing and Containing SARS-CoV-2

**DOI:** 10.1101/2021.09.01.458653

**Authors:** Xin Sun, Shaobo Yang, Amal A. Al-Dossary, Shana Broitman, Yun Ni, Mengdi Yang, Jiahe Li

**Affiliations:** Department of Bioengineering, Northeastern University, Boston, MA, United States, 02115; Department of Basic Sciences, Deanship of Preparatory Year and Supporting Studies, Imam Abdulrahman Bin Faisal University, Dammam, Saudi Arabia, 34212

**Author notes:** Corresponding author: Jiahe Li.

**Keywords:** COVID-19, SARS-CoV-2, nanobody, cellulose, cellulose binding protein

## Abstract

The highly transmissible severe acute respiratory syndrome coronavirus 2 (SARS-CoV-2) has infected more than 217 million people, claiming ~ 4.5 million lives to date. Although mandatory quarantines, lockdowns, and vaccinations help curb viral transmission, safe and effective preventative measures remain urgently needed. Here, we present a generic strategy for containing SARS-CoV-2 by cellulose materials. Specifically, we developed a bifunctional fusion protein consisting of a cellulose-binding domain and a nanobody (Nb) targeting the receptor-binding domain of SARS-CoV-2. The immobilization of the fusion proteins on cellulose substrates enhanced the capture efficiency of Nbs against SARS-CoV-2 pseudoviruses of the wildtype and the D614G variant, the latter of which has been shown to confer higher infectivity. Furthermore, the fusion protein was integrated into a customizable chromatography with highly porous cellulose for neutralizing virus from contaminated fluids in a continuous and cost-effective fashion. Taken together, our work leverages low-cost cellulose materials and recently developed Nbs to provide a complementary approach to addressing the pandemic.

**IMPORTANCE:** The ongoing efforts to address the COVID-19 pandemic center around the development of point-of-care diagnostics, preventative measures, and therapeutic strategies against COVID-19. In contrast to existing work, we have provided a complementary approach to target and contain SARS-CoV-2 from contaminated fluids and surfaces. Specifically, we present a generic strategy for the capture and containing of SARS-CoV-2 by cellulose-based substrates. This was archived by developing a bifunctional fusion protein consisting of both a cellulose-binding domain and a nanobody specific for the receptor-binding domain of SARS-CoV-2. As a proof-of-concept, our fusion protein-coated cellulose substrates exhibited enhanced capture efficiency against SARS-CoV-2 pseudovirus of both wildtype and the D614G mutant variants, the latter of which has been shown to confer higher infectivity. Furthermore, the fusion protein was integrated into a customizable chromatography with highly porous cellulose for neutralizing the virus from contaminated fluids in a highly continuous and cost-effective fashion.

## INTRODUCTION

Since the first documented coronavirus disease 2019 (COVID-19) case at the end of 2019 (1), the highly contagious severe acute respiratory syndrome coronavirus 2 (SARS-CoV-2) has resulted in at least 196 million positive cases and 4.2 million deaths (~2.17% fatality rate) in 219 countries and territories (as of July 28, 2021, source: https://covid19.who.int/). To contain the spread of SARS-CoV-2, non-pharmaceutical interventions were originally deployed, including the employment of masks, handwashing, and public measures such as city lockdowns, travel restrictions, and social distancing. However, the long-term adherence to these preventative measures has led to severe societal and economic crises (2). Importantly, the approval and administration of several SARS-CoV-2 vaccines worldwide has helped to mitigate the pandemic waves with an ever-increasing immunization population (3). Nevertheless, COVID-19 poses a continued threat because of constantly emerging SARS-CoV-2 variants and relatively long duration for herd immunity (4). Therefore, there is a great demand for effective, low-cost, and off-the-shelf agents to fast diagnose and decontaminate SARS-CoV-2 from body fluids and frequently contacted environmental surfaces (5).

SARS-CoV-2 belongs to the *β-coronavirus* genus of the *Coronaviridae* family, and shares the same subfamily *Orthocoronaviridae* with SARS-CoV, all of which lead to severe respiratory tract illness in humans (6). SARS-CoV-2 is a single-stranded RNA composed of 30 kb nucleotides, which encode four major structural proteins: the spike (S), the membrane (M), the envelope(E), and the nucleocapsid (N) (7). Viral infections rely upon cellular entry to utilize the host’s machinery for replicating viral copies that are then released by the host. The S protein facilitates the attachment of the virus to the host’s cellular receptors and promotes the fusion between host and viral membranes (8). In particular, the S protein contains the receptor-binding domain (RBD), which binds to the extracellular domain of the host receptor angiotensin-converting enzyme 2 (ACE2) for viral entry (9–12). Recent work demonstrates that SARS-CoV-2 targets the same functional host receptor, ACE2, as SARS-CoV (6,7,13). However, SARS-CoV-2’s RBD is ~10- to 20-fold higher than that of SARS-CoV in ACE2 binding (14). Owing to the key roles of SARS-CoV-2’s S protein or the subdomain RBD in the entry to host cells, the S protein or RBD has been extensively explored as a key target for the development of antiviral antibodies, among which nanobodies (Nbs) represent a unique class towards these efforts. Nbs are single-domain nanosized antibodies, which are derived from variable fragments of *Camelidae* (including camels and llamas) heavy chain-only antibodies (hcAbs) (15,16). Nbs offer a variety of advantages over other antibodies for diagnostic development: (i) nanometer size, (ii) high affinity and specificity, (iii) deep penetration in tissues, (iv) low immunogenicity, and (v) easy scalability for mass production (16,17). Because of these benefits, to date, several high-affinity neutralizing Nbs directed against SARS-CoV-2’s S protein and RBD domain have been identified, among which Ty1 has shown nanomolar binding affinity and effective neutralization through immunization in alpaca followed by the phage display (18–22). It was found that Ty1 specifically targets the RBD of SARS-CoV-2 with high affinity and directly blocks ACE2 engagement. Therefore, Ty1 can serve as a potential biologic for various diagnostic and therapeutic applications against COVID-19.

In this study, we repurposed the Nb Ty1 to detect and neutralize SARS-CoV-2 on surfaces and in biologically relevant fluids in a low-cost platform based on cellulose materials. Specifically, we designed a bifunctional fusion protein that comprises of a cellulose-binding domain (CBD) and Ty1 for cellulose immobilization and SARS-CoV-2 capturing, respectively. (23,24) The CBD originating from the *cipA* gene in the bacterium *C. thermocellum* has been shown to be resistant to heat denaturation (Tm ≥ 70 °C) due to the thermophilic nature of *C. thermocellum* (25). Nbs are generally easier to manufacture (e.g., *E. coli* fermentation) than conventional human immunoglobin (IgG) based antibodies, the latter of which require mammalian cell hosts for production (26). Additionally, use of CBD fusion proteins has been demonstrated for the biofunctionalization of cellulose substrates in applications including protein purification (27,28), textile manufacturing (29), and immunoassay development (30–33). Notably, the CBD domain can facilitate the absorption of CBD-containing fusion proteins to cellulose in molar quantities, which allows for an excess amount of immobilized proteins relative to the soluble target (30,34). As a result, our fusion technology is highly cost-effective and scalable to overcome various challenges posed by the pandemic, including but not limited to, disrupted supply chains, restricted deployment to remote areas, and mass production. As a proof-of-concept, we performed an immunoassay on cellulose-based filter paper for detecting SARS-CoV-2’s RBD using our bifunctional proteins. Furthermore, we developed a customized cellulose-based affinity chromatography to remove SARS-CoV-2 viral particles from biologically relevant fluids using pseudoviruses of both wildtype and the D614G variant, which may pave the way for decontaminating body fluids, including blood products (35,36). Given the modularity of our bifunctional protein platform and the ease of rapidly identifying target-specific Nbs through immunization and directed evolution, our work can potentially provide a framework to address other emerging infectious diseases with similar approaches.

## RESULTS

### Genetic fusion of an RBD-specific nanobody with a cellulose-binding domain

Our overall scheme for low-cost detection and neutralization of SARS-CoV-2 capitalizes on generating a bifunctional protein through the genetic fusion between the high-affinity Nb Ty1 targeting the RBD of SARS-CoV-2 and the cellulose-binding domain (CBD) (**Figure 1**). SARS-CoV-2 can be transmitted via airborne particles or through directly contacting contaminated surfaces (37). Therefore, considering that cellulose is prevalent in many materials, such as paper towels and the inner coating of face masks, we immobilized the fusion proteins to the surface of cellulose materials, such as filter paper, to enable viral detection or capturing of SARS-CoV-2 from the surface (**Figure 1e**). Importantly, due to the specific interaction between the CBD domain and cellulose, we reason that the bifunctional Ty1-CBD can be immobilized in a defined orientation that favors the interaction between the Nb of interest and the antigen, as compared to random immobilization. On the other hand, viral transmission and contamination through blood products present a common issue during the pandemic (36). Because blood products are susceptible to heat and chemical denaturation, it is desirable to devise a strategy that reduces and eliminates the viral load in blood products while maintaining the blood’s bioactivities (35). To assess whether our strategy can address these challenges, we integrated the bifunctional fusion protein Ty1-CBD into a customized cellulose affinity purification column to allow for a target-specific depletion in a continuous manner (**Figure 1f**).

**Figure 1:**
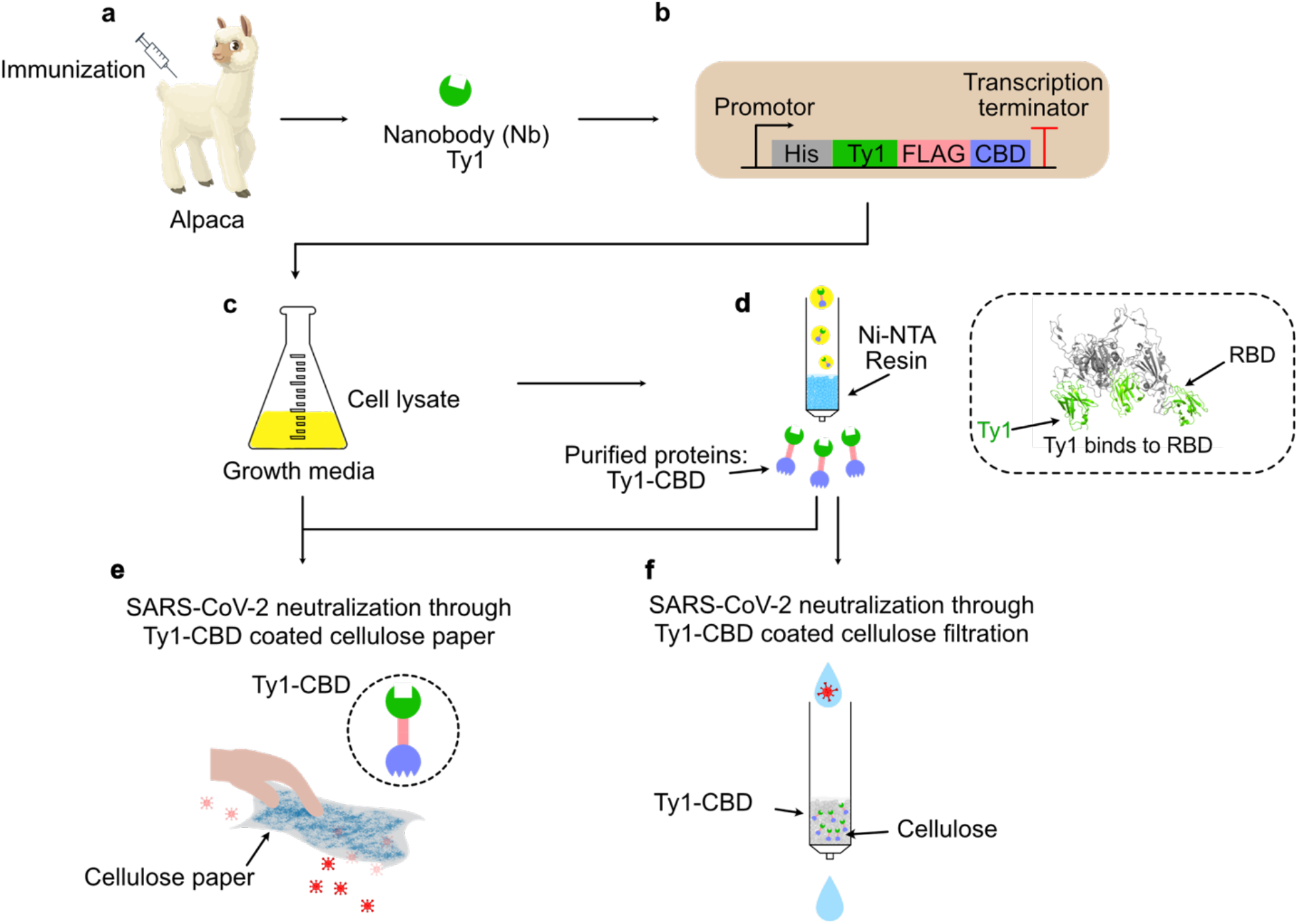
Development of a bifunctional fusion protein to enable cellulose immobilization and subsequent neutralization of severe acute respiratory syndrome coronavirus 2 (SARS-CoV-2). **a**) An alpaca-derived high-affinity nanobody (Nb), Ty1 for the receptor-binding domain (RBD) of SARS-CoV-2 was genetically fused with **b**) a cellulose-binding domain (CBD) isolated from a *C. thermocellum*. **c**) The fusion protein Ty1-CBD was recombinantly expressed in *E. coli*. **d**) The purification of CBD fusion proteins via nickel-nitrilotriacetic acid (Ni-NTA) or direct usage of *E. coli* cell lysate containing the CBD fusion protein for cellulose immobilization. The RBD of SARS-CoV-2 (in grey) bound with Nb Ty1 (in green) was adapted from the Protein Data Bank (PDB): 6ZXN. **e**) As SARS-CoV-2 is transmitted through surface contact, CBD fusion proteins or CBD-contained *E. coli* cell lysate were immobilized on the surface of cellulose materials, such as cellulose paper, for viral detection and capturing. **f**) Since SARS-CoV-2 can also be transmitted through blood products, we also customized Nb-dependent regenerated amorphous cellulose (RAC) materials to capture and deplete the virus from bodily fluids.

### Production and purification of the bifunctional protein Ty1-CBD

Compared to human IgG, Nbs can be produced in *E. coli* with high yield and purity (**Figure 1a**). Therefore, DNA encoding the fusion protein Ty1-CBD was first cloned in a standard expression vector for recombinant protein production in *E. coli*. Because the antigen-binding site of Nbs is closed to the N terminus, we placed the CBD at the C terminus of Ty1 to circumvent potential steric hindrance. Meanwhile, a short 6x histidine (His-tag) was attached to the N terminus for the metal affinity purification, while a FLAG epitope (DYKDDDDK) was inserted between Ty1 and the CBD, simultaneously serving as a hydrophilic flexible linkage and a tag for immunostaining (**Figure 1b**). The yield of fusion proteins was estimated to be ~50 mg per liter of bacterial culture in a shake-flask mode and was purified to high homogeneity as evidenced by denaturing SDS-PAGE and size exclusion chromatography (**Figure 1c**, **Figure 1d**, **Figure S1a**, and **Figure S1b**).

### The fusion protein is functionally active in cellulose binding and RBD detection

Next, we sought to evaluate whether the fusion proteins were capable of cellulose binding and Nb-specific target recognition. To this end, we first spotted purified fusion proteins on the surface of cellulose paper (**Figure 2a**). Upon air drying, the paper was stained with a rat antibody against the FLAG epitope in the fusion protein, followed by an anti-rat secondary antibody conjugated with horseradish peroxidase (HRP). A dark precipitate was visualized after incubation with an HRP substrate, 3,3’-Diaminobenzidine (DAB). To quantify the binding efficiency of the fusion protein to the cellulose paper, we immobilized serially diluted fusion proteins within a defined area on the Whatman filter paper followed by immunostaining with an anti-FLAG antibody. Indeed, the extent of protein immobilization correlated with the staining in a certain concentration range (**Figure 2b** and **Figure S1c**), from which we estimated that a surface area of 1 mm^2^ can be saturated by 500 ng (~0.02 nmol) of Ty1-CBD proteins in the Whatman filter paper.

**Figure 2:**
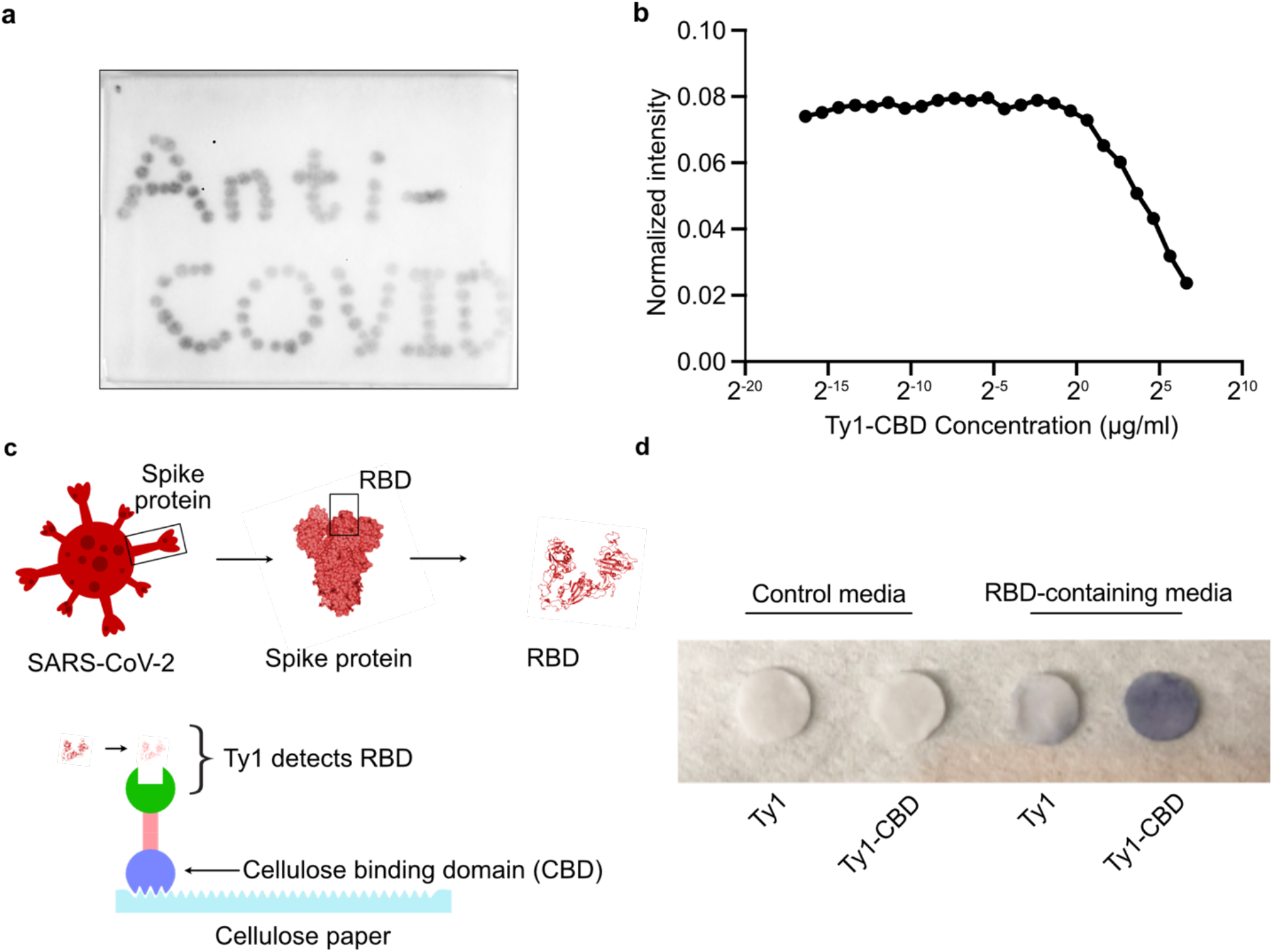
The fusion protein maintains its activities in binding cellulose and the RBD domain of SARS-CoV-2. **a**) Detection of immobilized Ty1-CBD on a cellulose paper. Ty1-CBD was first spotted on a piece of cellulose paper. Upon air drying, the paper was incubated with a rat antibody against the FLAG epitope (DYKDDDDK), followed by an anti-rat secondary antibody conjugated with HRP. The dark precipitate “Anti-COVID” was visualized after incubation with 3,3’-Diaminobenzidine (DAB). **b**) Quantification of maximal protein absorption on Waterman filter paper. 10 μL of serially diluted fusion protein solutions were applied to the filter paper, followed by immunoblotting with anti-FLAG directly on the filter paper. Based on the normalized unit intensity quantified by ImageJ, protein abundance increased with concentration. We estimate that 500 ng of Ty1-CBD binds to 1 mm^2^ of cellulose paper in saturation status. **c**) Schematic of an immunoassay to evaluate the function of the fusion protein. Ty1-CBD fusion proteins were immobilized on cellulose paper and then submerged in culture media containing RBD-Fc (~100 ng/ml) as a proxy for actual SARS-CoV-2. The capturing capability was confirmed by anti-human Fc-HRP and the DAB substrate. The structure of the RBD was adapted from PDB: 6ZXN. **d**) Testing the ability of protein-coated cellulose paper discs in capturing RBD-containing media. Representative discs were prepared by a 6-mm biopsy punch and then coated with *E. coli* lysates containing indicated recombinant fusion proteins. The functionalized discs were incubated with RBD-containing or control (no RBD) media. The intensity of dark staining was strongest from the combination of Ty1-CBD-coated disc and RBD-containing media (~100 ng/ml).

Having validated the high binding capacity of purified Ty1-CBD to filter paper, we speculated that because the CBD itself can act as a natural affinity ligand to cellulose, we could directly immobilize *E. coli* cell lysate containing the CBD fusion proteins to filter paper, followed by extensive washing to remove nonspecific proteins. This circumvents the need to purify desired proteins beforehand, which is labor intensive and impractical when it comes to the large-scale manufacturing of functionalized cellulose materials (**Figure 2c**). To this end, we first incubated *E. coli* cell lysate with filter discs and removed nonspecific proteins via washing. Then we subjected the functionalized filter discs to cell culture media containing secreted recombinant RBD as a proxy for actual SARS-CoV-2. As shown in **Figure 2d**, Ty1-CBD-coated filter paper was able to capture SARS-CoV-2’s RBD as evidenced by the intense dark staining reflecting the detection of RBD. Of note, filter discs precoated with Ty1 alone exhibited light staining, likely due to nonspecific but weak absorption of Ty1 to cellulose. In comparison, in the control media without the RBD, neither Ty1- nor Ty1-CBD-coatd discs displayed the dark staining. Our findings here indicate that the CBD domain promoted the immobilization of Ty1-CBD on cellulose substrates while Ty1 remained able to specifically recognize the target.

### Immobilization of Ty1-CBD on filter paper increases the capture efficiency of Ty1 against SARS-CoV-2 pseudovirus

The stoichiometry and kinetics of a target-binding interaction can be favorably influenced by three general approaches: (i) increasing the molar abundance and the soluble antigen concentration, (ii) improving the binding interaction affinity under relevant assay conditions, and (iii) raising the capturing reagents (e.g. antibodies) abundance and concentration through surface immobilization according to the law of mass action (38,39). Since it is not practical to raise the concentration of antigens or the affinity of already optimized antibodies, here we sought to explore the third strategy of increasing the surface densities of Ty1-CBD via immobilization on cellulose paper. To do so, we evaluated the capability of Ty1-CBD fusion proteins in capturing SARS-CoV-2 mimics (referred to as pseudovirus in this work). One of the gold standards for SARS-CoV-2-related studies is to use non-replicative lentivirus pseudotyped with the S protein derived from SARS-CoV-2 in conjunction with mammalian cells engineered to express human ACE2 (hACE2) (40) (**Figure 3a**). Since emerging SARS-CoV-2 variants with higher transmission rates bear mutations in the S protein, one advantage of the pseudovirus system is that it can rapidly evaluate intervention approaches against different spike variants. Using this system, we compared the original wildtype (WT) S spike to the “D614G” mutant, in which the 614^th^ aspartate is converted to glycine in the S protein of SASR-CoV-2. Notably, epidemiology and molecular biology studies have demonstrated that the D614G mutant confers higher transmission and worse symptoms in humans (41). Therefore, it is of particular interest to assess our fusion protein strategy in the context of both the wildtype and the D614G variant. As shown in **Figure 3b** and **3c**, after HEK293T-hACE2 cells were transduced with wildtype or D614G pseudotyped lentivirus carrying a green fluorescence protein (GFP) reporter, ~50% of cells were GFP positive with D614G pseudovirus exhibiting a higher transduction efficacy than that of the wildtype. These findings agreed with the increased infectivity by the D614G mutation (42,43). In comparison, transduction of the parental HEK293T cell line (lacking hACE2 expression) with the same SAR-CoV-2 pseudovirus did not result in GFP expression, which validated an ACE2-dependent infection by SAR-CoV-2 (44).

**Figure 3:**
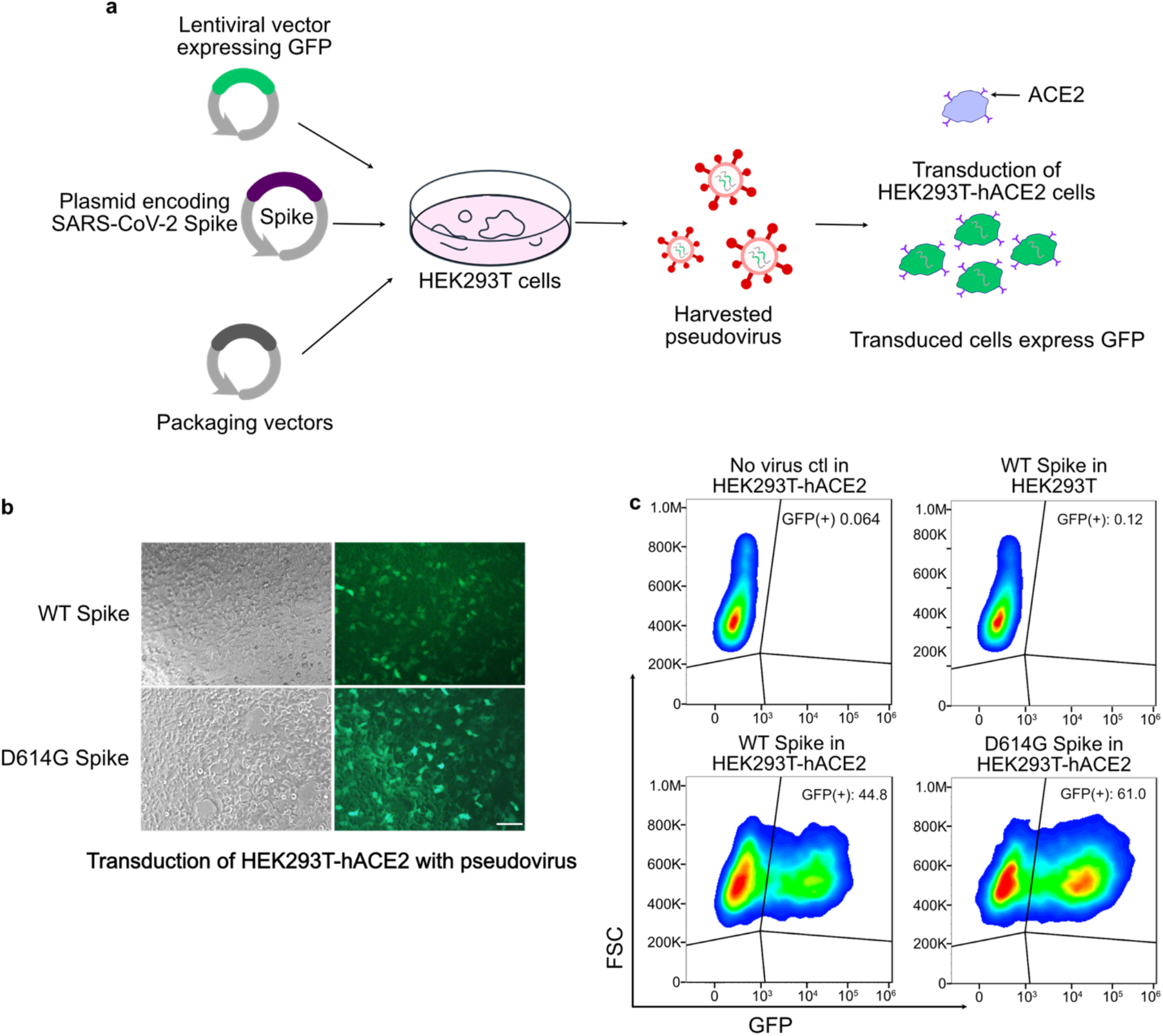
Generation of wildtype and D614G pseudoviruses for neutralization assays by Ty1-CBD-functionalized cellulose. **a**) Schematic overview of the pseudotyped virus production: HEK293T cells were transfected with a lentiviral vector expressing a green fluorescent protein (GFP), a plasmid encoding SARS-CoV-2 spike, and packaging vectors. The transfected cells produced lentiviral particles pseudotyped with the S protein of SARS-CoV-2, and the pseudovirus can transduce HEK293T expressing human angiotensin-converting enzyme 2 (hACE2) to express GFP. **b**) Microscope images showing that the HEK293T-hACE2 cells expressed GFP after transduction with lentivirus pseudotyped with the wildtype (WT) SARS-CoV-2 spike protein or the D614G variant. Scale bar = 100 μm. **c**) Representative flow cytometric analysis evaluating the transduction efficiency of SARS-CoV-2 WT and D614G pseudoviruses compared with two negative control groups: HEK293T-hACE2 without any transduction and HEK293T transduced with SARS-CoV-2 WT pseudotyped lentivirus. Results are representative of three independent experiments.

It is worth noting that the levels of SARS-CoV-2 in COVID-19 patients range from 10^4^ to 10^9^ copies per ml depending on the type of bodily fluids and degree of the symptoms (45,46). Meanwhile, we calculated the titers of wildtype or D614G pseudotyped lentivirus, and estimated that ~10^5^ viral particle particles per ml were present in the culture media. Therefore, to demonstrate the capability of capturing SARS-CoV-2 at the lower end of the viral titer range for SARS-CoV-2-containing fluids, we further diluted the media to contain approximately ~10^4^ pseudovirus copies per ml, and quantified the neutralization efficacy of Ty1-CBD-immobilized cellulose paper (**Figure 4a**). In addition, filter paper alone or filter paper coated with Ty1 but lacking the CBD module served as negative controls. The neutralization efficiency of Ty1-CBD-immobilized filter paper was calculated by dividing the titer of media treated with Ty1-CBD-immobilized filter paper or other control groups by the initial viral titer (i.e., without any treatment). Indeed, using media containing wildtype or D614G pseudoviruses (**Figure 4b**), Ty1-CBD-immobilized filter paper resulted in ~2-fold increase of the neutralization efficacy compared to that of filter paper only, and ~1.5-fold enhancement over filter paper pre-coated with Ty1 alone. Moreover, filter paper pre-coated with Ty1-CBD or Ty1 displayed ~1.65-fold improvement in neutralizing pseudovirus from the media over free proteins (Ty1-CBD or Ty1) at equal concentrations, which indicated that the surface immobilization itself can facilitate target recognition.

**Figure 4:**
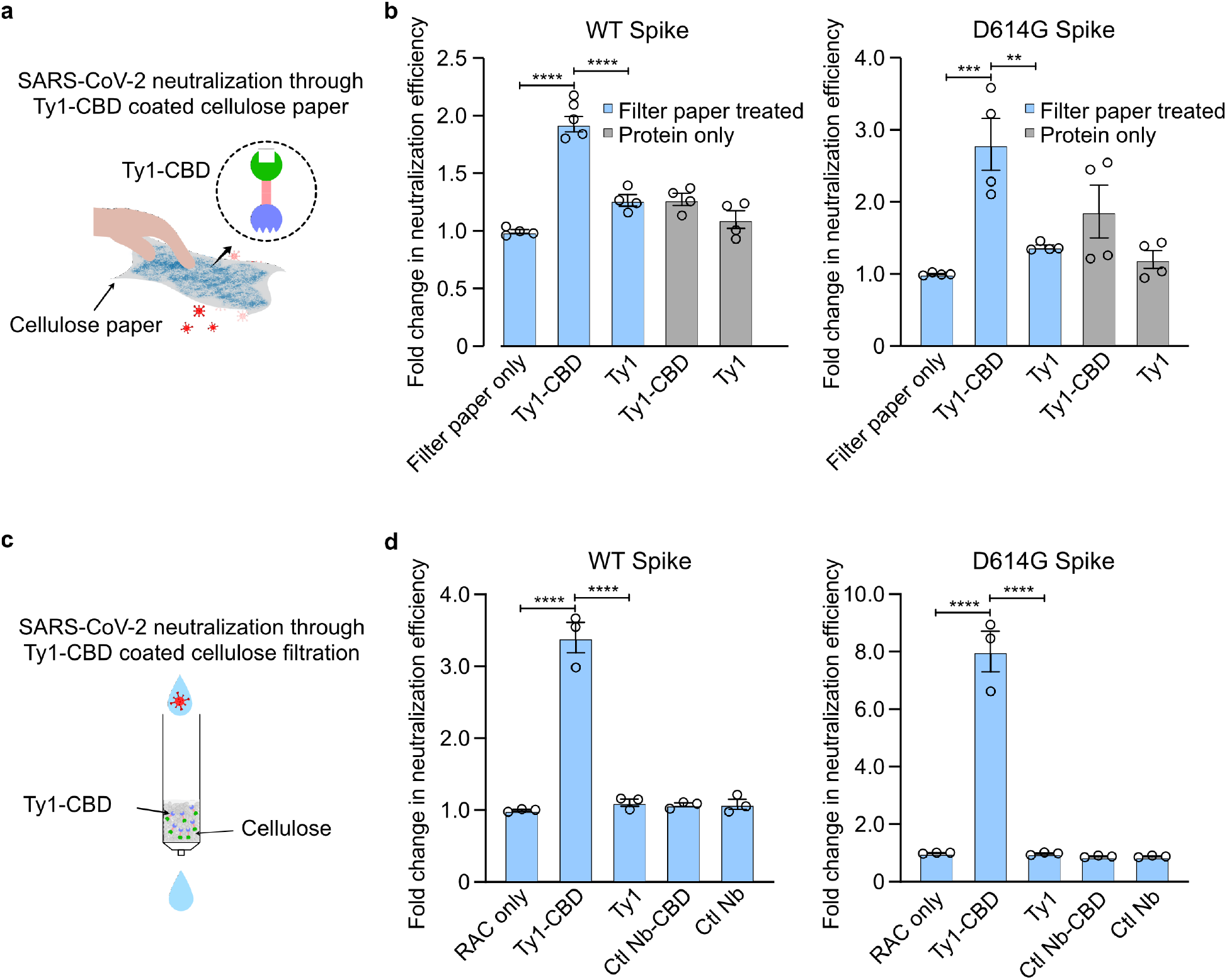
SARS-CoV-2 pseudovirus capture by Ty-CBD-immobilized cellulose in two different formats: **a**) Schematic of increasing surface densities of Ty1-CBD through protein immobilization on cellulose materials for SARS-CoV-2 neutralization. **b**) Increased neutralization efficacy of pseudovirus through protein immobilization on cellulose paper over free proteins. After incubating the pseudovirus with 200 μL of 10 μg/ml fusion protein Ty1-CBD or Ty1 (negative control) immobilized on cellulose paper or free protein with equal concentrations, the titers of wildtype (WT) and D614G pseudoviruses were quantified by transducing HEK293T-hACE2 cells with the remaining viruses in the supernatant. Fold changes from each treatment group were normalized to that of filter paper only. **c**) A Ty1-CBD-functionalized RAC column to capture the antigen of interest in a continuous fashion. **d**) Neutralization efficacy of Ty1-CBD-functionalized RAC. The flow through samples from functionalized RAC columns were used to transduce HEK293T-hACE2 cells to quantify viral titers for WT and D614G SARS-CoV-2 pseudoviruses, respectively. Fold changes from each treatment group were normalized to that of RAC only. Graphs are expressed as mean ± SEM (*n* = 4) in **b** and as mean ± SEM (*n* = 3) in **d**. Statistical analysis was performed by one-way analysis of variance (ANOVA) according to the following scale: \*\**P* < 0.01, \*\*\**P* < 0.001, and \*\*\*\**P* < 0.0001.

### Integration of the bifunctional protein with an amorphous cellulose column further enhanced the capture efficiency

Having validated the increased capture efficiency of fusion proteins immobilized on filter paper, we further sought to enhance the neutralization efficiency by incorporating the fusion proteins into regenerated amorphous cellulose (RAC). Because RAC has been shown to exhibit higher surface area per unit mass than filter paper (47), we reasoned that using RAC can increase the immobilization density of Ty1-CBD on cellulose, which in turn improves the rate and the degree of target capture based on the theoretical modeling by others (23,24). Moreover, this strategy can potentially expand the utility of Ty1-CBD by packing Ty1-CBD-functionalized RAC in a column, which allows for an affinity chromatography system to reduce viral load from contaminated fluids (e.g., blood and saliva) in a continuous mode. To test our hypotheses, we filled a 1 mL gravity-based column with 0.1 mL (~50 mg dry weight) of RAC, which was subsequently saturated with purified Ty1-CBD proteins or *E. coli* lysate containing the fusion proteins (**Figure 4c**). After culture media containing wildtype or D614 pseudoviruses were passed through the functionalized column by gravity, viral titers were determined for different flow through samples. Compared to RAC alone, Ty1-CBD-immobilized columns increased the neutralization efficacy of wildtype and D614G pseudoviruses by ~ 3.5 times and ~8 times, respectively. In contrast, RAC columns carrying an irrelevant Nb (caffeine specific) fused with CBD or Ty1 alone failed to further enhance the degree of neutralization compared to that of RAC alone (**Figure 4d**). Taken together, we demonstrated that the Ty1-CBD fusion protein can be integrated into an RAC column to markedly increase the neutralization efficiency of SARS-CoV-2 pseudovirus in a highly specific and continuous fashion.

## DISCUSSION

In comparison to others’ efforts in the development of cost-effective point-of-care (POC) diagnostics, preventative measures, and therapeutic strategies against COVID-19, we have developed a bifunctional fusion protein technology that features SARS-CoV-2 capture from cellulose substrates. Since cellulose represents the most abundant and commonly used biopolymer, our approach holds the promise to enable cellulose-based POC diagnostics, functionalized face masks to reduce airborne virus transmission, and customized affinity columns to decontaminate fluids containing SARS-CoV-2 (48,49). Notably, a similar strategy has been proposed for SARS-CoV-2 detection through cellulose filter paper immobilized with CBD fusion proteins (23,24). In their approach, the nucleocapsid protein was fused with the CBD and the detection of SARS-CoV-2 required an antibody to capture viral particles in a “sandwich” format. In comparison, we have developed a single agent by directly linking CBD with a Nb specific for SARS-CoV-2, which may be more convenient and cost effective. Additionally, we have demonstrated that the bifunctional fusion proteins can serve as an affinity agent in the customized cellulose column to deplete virus from fluids. Future work can compare these two approaches in terms of capture efficiency and specificity. In addition, we showed that ~ 50 mg of Ty1-CBD was produced from one-liter bacterial culture in a shake-flask mode, which in theory could functionalize cellulose paper with a surface area of 0.1 m^2^ based on our titration experiments. Future work can exploit advanced fermentation technologies to improve the production yield of the fusion proteins.

While this study only focused on Ty1, a recently developed Nb against the spike protein of SARS-CoV-2 (18), the conceptual framework can be easily adapted to target other emerging viruses by substituting the Nb module with other target-specific Nbs. Moreover, since many other SARS-CoV-2-specific Nbs have been identified through immunization and the phage display to target different epitopes for SARS-CoV-2, future work may investigate a combination of CBD fusion proteins comprising different Nbs in a multivalent manner to enhance the capture efficiency (50). Despite the promises demonstrated in this study, one limitation is that only pseudovirus-containing culture media were used to characterize the fusion proteins for proof-of-concept. Therefore, it is necessary to further evaluate our approach in real specimens such as blood from COVID-19 patients in the near future.

## Materials and Methods

### Reagents and chemicals

Tween-20, Triton X-100 (TX-100) and Triton X-114 (TX-114) were obtained from Sigma-Aldrich (St Louis, MO). Strep-tag and Strep-Tactin XT were purchased from IBA Lifesciences (Gottingen, Germany). Detergent compatible (DC) protein assay kit was bought from Bio-Rad Laboratories (Hercules, CA). Anti-FLAG epitope (DYKDDDDK, catalog# 637301) and horseradish peroxidase (HRP) Donkey anti-human IgG antibody (catalog #410902) were purchased from Biolegend (San Diego, CA). The secondary antibody anti-rat IgG HRP (catalog# 7077) was bought from Cell Signaling Technology (CST, Danvers, MA). All other reagents and chemicals, including nickel-nitrilotriacetic acid (Ni-NTA) agarose and Pierce Rapid Gold BCA Protein Assay Kit were purchased from Fisher Scientific International Inc., (Hampton, NH), and were of highest purity or analytical grade commercially available.

### Cell lines

HEK293T cells expressing human angiotensin I-converting enzyme 2 (HEK293T-hACE2) were kindly provided by Dr. Jesse Bloom (Fred Hutchinson Cancer Research Center, Seattle, USA) (51). Lenti-X 293T cell line was purchased from Takara Bio USA Inc. (San Jose, CA). These cell lines were maintained in complete Dulbecco’s modified Eagle’s medium (DMEM) (Corning, NY) supplemented with 10% fetal bovine serum (FBS; Corning, NY) and 100 U/ml penicillin-streptomycin (Corning, NY) at 37 °C in humidified incubator with 5% carbon dioxide (CO_2_). Cells at passages 2-10 were used for the experiments.

### Plasmids construction, protein expression and purification

Ty1 variants, including Ty1-CBD and control protein Ty1 without CBD module, were cloned into pSH200 vector (a generous gift from Prof. Xiling Shen at Duke University) containing 6 x Histidine tag (His-tag), between *BamHI* and *XbaI* sites. Both plasmids were validated by sequencing before expression. Ty1-CBD and Ty1 were expressed and produced in the same manner as previously described (Yang et al., 2020; Sun et al., 2021). Produced protein was sequentially purified by affinity chromatography using Ni-NTA agarose beads and fast protein liquid chromatography (FPLC, NGC Quest 10 Chromatography system, Biorad, Hercules, CA). Protein fractions detected at λ = 280 were collected. Collected protein fraction were quantified by detergent compatible (DC) protein assay according to the manufacturer’s instructions and purities were verified by sodium dodecyl sulfate-polyacrylamide gel electrophoresis (SDS-PAGE). Validated protein was aliquoted and kept at −80 °C with 50% glycerol at all times until future use.

### Cellulose paper-based immunoblotting

To validate the binding ability of Ty1-CBD, purified Ty1-CBD was spotted on cellulose paper and dried at room temperature for ~2 min. Cellulose paper with dried Ty1-CBD were incubated with 5 ml of 5% nonfat milk in Tris-buffered saline (TBS) for 30 min to block non-specific binding sites. After blocking, the cellulose paper was then incubated with anti-FLAG epitope (DYKDDDDK), which was diluted at 1:2000 in 3 ml TBS plus 5% nonfat milk, overnight at 4°C. The paper was washed three times with 5ml 1xTBS containing 0.05% Tween-20 (TBST), with 15 min per wash cycle. The paper was incubated with HRP anti-rat IgG (1: 2000) for 1 hr at room temperature. After washing with 1xTBS with 5 ml 0.05% Tween-20, premixed Pierce™ 3,3’diaminobenzidine (DAB) substrate was directly added on the cellulose paper. The reaction was terminated with water after dark spots appeared.

### Pseudovirus production

HDM-SARS2-Spike-delta21 and HDM-SARS2-Spike-del21-614G encoding SARS-CoV-2 Spike with 21 amino acid C-terminal deletion for lentiviral pseudo-typing were purchased from Addgene (Watertown, MA) with serial numbers of Addgene #155130 and Addgene#158762, respectively. Both plasmids were isolated following manufacturer’s instructions and quantified through Nanodrop. 24 hr prior to transfection, Lenti-X 293T cells in logarithmic growth phase were trypsinized, and the cell density was adjusted to 1.0×10^6^ cells/mL with complete DMEM medium. The cells were reseeded into a 10 cm cell culture dishes to reach 70% confluency on the day of transfection. The plasmid mixture was prepared according to **Table 1** and **Table 2**. After inverting the plasmid mixture for 5-8 times, and kept at room temperature for 30 min, the plasmid mixture was gently added to the Lenti-X 293T cells (51). After 48 hr and 72 hr of transfection, the pseudovirus was collected and centrifuged at 1000xg at 4 °C for 20 min to remove the debris. The cell culture medium was replaced with fresh complete DMEM medium once the pseudovirus was collected. Aliquots of the harvested virus were stored at 4 °C for immediate use or frozen at −80 °C for future use.

**Table 1.** Compositions of the plasmid mixtures for generating SARS-CoV-2 pseudovirus
(wildtype or D614G).

| Composition | Amount |
| --- | --- |
| HDM-SARS-2-Spike-delta21 (or D614G) | 2.5 µg |
| pFuw-Ubc-GFP | 10 µg |
| psPAX2 | 7.5 µg |
| TransIT-X2 2000 | 30 µl |
| Opti-MEM | 1000 µL |

**Table 2:** Compositions of the plasmid mixture for lentivirus generation as a positive control.

| Composition | Amount |
| --- | --- |
| pMD2.G | 2.5 µg |
| pFuw-Ubc-GFP | 10 µg |
| psPAX2 | 7.5 µg |
| TransIT-X2 2000 | 30 µl |
| Opti-MEM | 1000 µL |

### Quantification of viral titers by flow cytometry

HEK293T-hACE2 cells were seeded 24 hr before the pseudovirus transduction assay in 96-well plates (2×10^4^ cells/well in a volume of 100 μl complete DMEM medium). On the co-culture day, the medium was removed and 200 μl of prewarmed pseudovirus was added to the cells. Polybrene (Sigma Aldrich) was added into cultured HEK293T-hACE2 cells to a final concentration of 8 μg/ml, to facilitate lentiviral infection through minimizing charge-repulsion between virus and cells. After transduction for 48 hr, cells were collected through trypsinization and transferred to a 96 well V-bottom plate with complete culture DMEM medium for neutralization. Cells were pelleted at 300 xg for 3 min and washed twice with phosphate-buffered saline (PBS, 137 mM NaCl, 10 mM phosphate, 2.7 mM KCl; pH 7.4). After the final wash, the cells were resuspended in 200 μL FACS buffer (5% FBS, 2mM EDTA, 0.1% sodium azide in PBS) for flow cytometric analysis in Attune™ NxT Flow Cytometer (Thermofisher) and data were analyzed by FlowJo (Franklin Lakes, NJ). Using a well that has 1%-20% of GFP-positive cells, the titer was calculated according to the formula, titer = {(F × Cn) /V}× DF, F: The frequency of GFP-positive cells determined by flow cytometry; Cn: The total number of target cells infected; V: The volume of the inoculum. DF: The virus dilution factor. For all experiments triplicate samples were analyzed, data are representative of two or more experiments, and the standard error of the mean (SEM) is shown.

### Preparation of the regenerated amorphous cellulose (RAC) column

RAC was produced referring to RAC-based affinity protein purification as previously described (47). The RAC column was first equilibrated with 1xPBS and proteins of interest were added to the RAC column until the column was saturated by quantifying the amount of fusion protein in flow-through fractions and comparing it to the original supernatant via SDS-PAGE. The saturated RAC was washed with 10 column volumes of 1xPBS followed by loading with 400 μl complete DMEM medium. 400 μL of pseudovirus were flowed through the saturated RAC and collected for future use.

### Preparation of Ty1-CBD functionalized cellulose paper

For capturing capability assay, cellulose filter paper discs were fitted into a 96-well plate or 1.5 ml microtube followed by blocking with 1% bovine serum albumin (BSA) in 1 x TBS for 1 hr. After aspirating the blocking buffer, 50-100 μl purified proteins with concentrations of 100 μg/ml, 10μg/ml and 1μg/ml in 1 x TBS or 400 μl protein lysate of interest were applied directly to the coated cellulose paper. After 1 hr incubation at room temperature with slow shaking, purified proteins or lysates were removed from 96 well plates or 1.5 ml microtube, followed by 3 times wash with 1x TBS, 10 min in between. Cellulose paper was blocked with 5% nonfat milk in TBS at room temperature for 15 – 30 min. 50 – 100 μL media with spike expression were applied to the cellulose paper and incubated at room temperature for 1 hr. The treated cellulose paper was washed with TBST, 4 times, with 5 -10 min each. The washed cellulose paper was incubated with HRP Donkey anti-human IgG antibody diluted in 1 x TBS supplemented with 5% milk with antibody dilution of 1:2000. After 1 hr, the cellulose paper was washed with TBST for 4 times with 5 – 10 each. Premixed DAB substrate was directly added on cellulose paper after removing the TBST. The reaction was terminated with distilled water till the color development.

For preparing the cellulose filter paper discs to evaluate the neutralization efficiency against the pseudovirus, cellulose filter paper discs were fitted into a 96-well or 48-well plate followed by blocking with 1% BSA in 1 x TBS for 1 hr. After aspirating the blocking buffer, 50 - 100 μl of 10 μg/ml purified fusion proteins in 1 x TBS were applied directly to the coated cellulose paper. After 1 hr incubation at room temperature with slow shaking, diluted purified proteins were aspirated from 96-well or 48-well plates, followed by two times wash with 1x TBS, 10 min in between. 100-400 μL of pseudoviruses were added into each well with cellulose paper. After 1 hr incubation, the cellulose paper treated pseudovirus was gently transferred to transfect HEK293T-hACE2 cells with 80% confluency, followed by adding polybrene to a final concentration of 8 μg/ml. 24 hr post-transduction, HEK293T-hACE2 cells were split in a ratio of 1:3. 48 hr post transfection, cells were collected at 300 xg for 3 min followed by resuspending them in 200 μL FACS buffer. Samples were loaded to Attune flow cytometry followed by analysis through FlowJo. After gating the HEK293T-hACE2 cells without pseudovirus (negative control) with less than 2% GFP positive, the transduction efficiency for experimental group is calculated by GFP^+^ / [GFP^+^+GFP^−^]. For all experiments triplicate samples were analyzed, data are representative of two or more experiments, and the standard error of the mean (SEM) is shown.

### Statistical analysis

Statistical significance was evaluated using one-way ANOVA followed by Tukey post hoc test using GraphPad PRISM (San Diego, CA, USA). *P* values < 0.05 were considered statistically significant. Statistical significance of our interest is indicated in all figures according to the following scale: *P<0.05, **P<0.01, ***P<0.001, and ****P<0.0001. All graphs are expressed as the means ± SEM.

## Acknowledgements

This work was supported by the Northeastern University COVID19 Rapid Seed Grant (JL), Peer Reviewed Medical Research Program from the Department of Defense’s Congressionally Directed Medical Research Programs (JL). We extend our appreciation to the Deputyship for Research & Innovation, Ministry of Education in Saudi Arabia (JL and AAA) for funding this research work. We would like to express our gratitude to Professor Sara Rouhanifard at Northeastern University’s Department of Bioengineering for generously sharing her lab’s fluorescent microscope; Professor Ke Zhang at Northeastern University’s Department of Chemistry for sharing his lab’s equipment.

## AUTHOR CONTRIBUTIONS

J.L. designed the study. X. S., S. Y., Y.N., M.Y., performed the experiments. J.L., X.S., S.Y., A.A., S.B., Y.N., M.Y., analyzed the data. J.L., X.S., and A.A. wrote the manuscript.

